# Interpreting biologically informed neural networks for enhanced biomarker discovery and pathway analysis

**DOI:** 10.1101/2023.02.16.528807

**Authors:** Erik Hartman, Aaron Scott, Lars Malmström, Johan Malmström

## Abstract

The advent of novel methods in mass spectrometry-based proteomics allows for the identification of biomarkers and biological pathways which are crucial for the understanding of complex diseases. However, contemporary analytical methods often omit essential information, such as protein abundance and protein co-regulation, and therefore miss crucial relationships in the data. Here, we introduce a generalized workflow that incorporates proteins, their abundances, and associated pathways into a deep learning-based methodology to improve biomarker identification and pathway analysis through the creation and interpretation of biologically informed neural networks (BINNs). We successfully employ BINNs to differentiate between two subphenotypes of septic acute kidney injury (AKI) and COVID-19 from the plasma proteome and utilize feature attribution-methods to introspect the networks to identify which proteins and pathways are important for distinguishing between subphenotypes. Compared to existing methods, BINNs achieved the highest predictive accuracy and revealed that metabolic processes were key to differentiating between septic AKI subphenotypes, while the immune system was more important to the classification of COVID-19 subphenotypes. The methodology behind creating, interpreting, and visualizing BINNs were implemented in a free and open source Python-package: https://github.com/InfectionMedicineProteomics/BINN.

## 1 Introduction

The continuous technological advancements in mass spectrometry based proteomics has the enabled the quantification of hundreds to thousands of proteins in clinical samples extending its reach in biomedical and clinical research [1, 2]. The increasing ability to rapidly analyze a large number of clinical samples provides new opportunities to profile complex biological systems and bridge the gap between translational and clinical research through the investigation of disease mechanisms and the identification of biomarkers. These advances are of interest for many disease areas, such as the study of infectious diseases where the identification of distinct clinical and molecular subphenotypes may impact the development of new treatment regimes. Subphenotypes are typically identified using clinical parameters based on the presented severity of different symptoms of the disease and are difficult to distinguish. Previous work has proposed clinical subphenotypes for COVID-19 [3, 4, 5, 6] and sepsis [7, 8, 9], but the development of targeted treatments for the different subphenotypes remains challenging as the underpinning molecular mechanisms are poorly characterized. To understand these molecular mechanisms, it is therefore critical to analyze the proteins and associated biological pathways of a disease to support the development of precision treatments and provide the best patient care possible.

Currently, a common strategy to identify candidate diagnostic and prognostic biomarkers are based on significantly differentially expressed (DE) proteins between subphenotypes. Substantial research has been conducted on how to optimize DE detection algorithms [10, 11, 12, 13, 14], but the process of selecting proteins for further investigation remains unstandardized. In most cases, proteins that pass a *p-value* and fold-change threshold are considered the most informative, but these thresholds are rule based and potentially eliminate important biological signal. To understand the systemic impact of DE proteins, it is also pertinent to identify which pathways are enriched based on the difference in abundance of the DE proteins. Several tools and databases have been developed to automate this process and to select the most significant pathways based on the proteins that have been identified in DE analysis [15, 16, 17]. Commonly, the significance of a pathway is determined by counting the number of DE proteins that are connected to the pathway in a database and calculating a *p-value* based on these connections. This type of analysis typically omits crucial information such as protein abundance, protein co-expression, and pathway co-regulation, and selects the most interesting pathways using *p-value* cut-offs.

To mitigate these limitations, increasing efforts have been directed towards incorporating machine learning methods into proteomics workflows to improve the study of disease mechanisms and biomarker discovery [18, 19, 20]. Recent advances in the field of machine learning have allowed deep neural networks to thrive in domains of high dimensionality where complex networks can learn representations of features without the need for feature selection algorithms [21]. However, complex machine learning models, such as deep neural networks, suffer from a lack of interpretability, and although they provide greater predictive power than their more interpretable linear counterparts, this questions the utility of such methods. Research in the field of explainable artificial intelligence (xAI) has resulted in methods which allow for the interpretation of complex models by calculating the importance of each feature to the output of the model [22, 23, 24]. To further improve interpretability, biologically informed neural networks (BINNs) establish connections between their layers based on biological processes [25, 26] and thus generalize to unseen data more effectively [27].

Here, we demonstrate the utility of BINNs to develop highly accurate predictive models that enhance blood-based proteomics biomarker identification while providing greater insight into the underlying biology of a system. We apply our method to analyze proteomic differences in blood plasma between subphenotypes in sepsis induced acute kidney injury (AKI) and COVID-19. The two subphenotypes of septic AKI were previously established using clinical and molecular parameters using latent class analysis [28]. Similarly, the COVID-19 subphenotypes were based on varying levels of severity, where patients in need of assisted mechanical ventilation comprise the more severe subphenotype [29]. The annotations from these predictive models were used to select potential diagnostic biomarker panels to augment the parameters used to stratify the subphenotypes from the above datasets while providing a molecular explanation for their physical manifestation. We also demonstrate how BINNs can be used for intelligent pathway analysis to extract the most important pathways in a biological system. Overall, the inherent interpretability of BINNs lend to their potential to investigate complex biological systems in a more comprehensive manner and to enhance the potential of biomarker discovery in mass spectrometry-based proteomics. A generalizable and user friendly software package for the creation and analysis of annotated sparse BINNs is open source and freely available at https://github.com/InfectionMedicineProteomics/BINN.

## 2 Results

Currently, common proteomics-based biomarker identification and biological pathway analyses are based on arbitrary thresholds which omits important relationships in datasets, and therefore lack the comprehensiveness which is required when analyzing complex biological systems. Here, we apply a deep learning-based methodology which utilizes the Reactome pathway database [16] to incorporate biological relationships in a biologically informed neural network (BINN), allowing for a unified analysis of biomarkers, biological pathways, and processes. The Reactome database contains information about relationships of biological entities, and its underlying graph is manipulated to fit a sequential neural network-like structure, resulting in a sparse architecture where nodes are annotated with a protein, biological pathway, or biological process. We create and employ BINNs on two proteomic datasets, distinguishing between two subphenotypes of septic akute kidney injury (AKI) and COVID-19. The BINNs are fed the proteomic content of samples and thereby trained to classify the subphenotypes, whereafter they are interpreted using Shapley Additive Explanations (SHAP) [22], eventually allowing for the identification of important proteins and pathways (figure 1).

**Figure 1:**
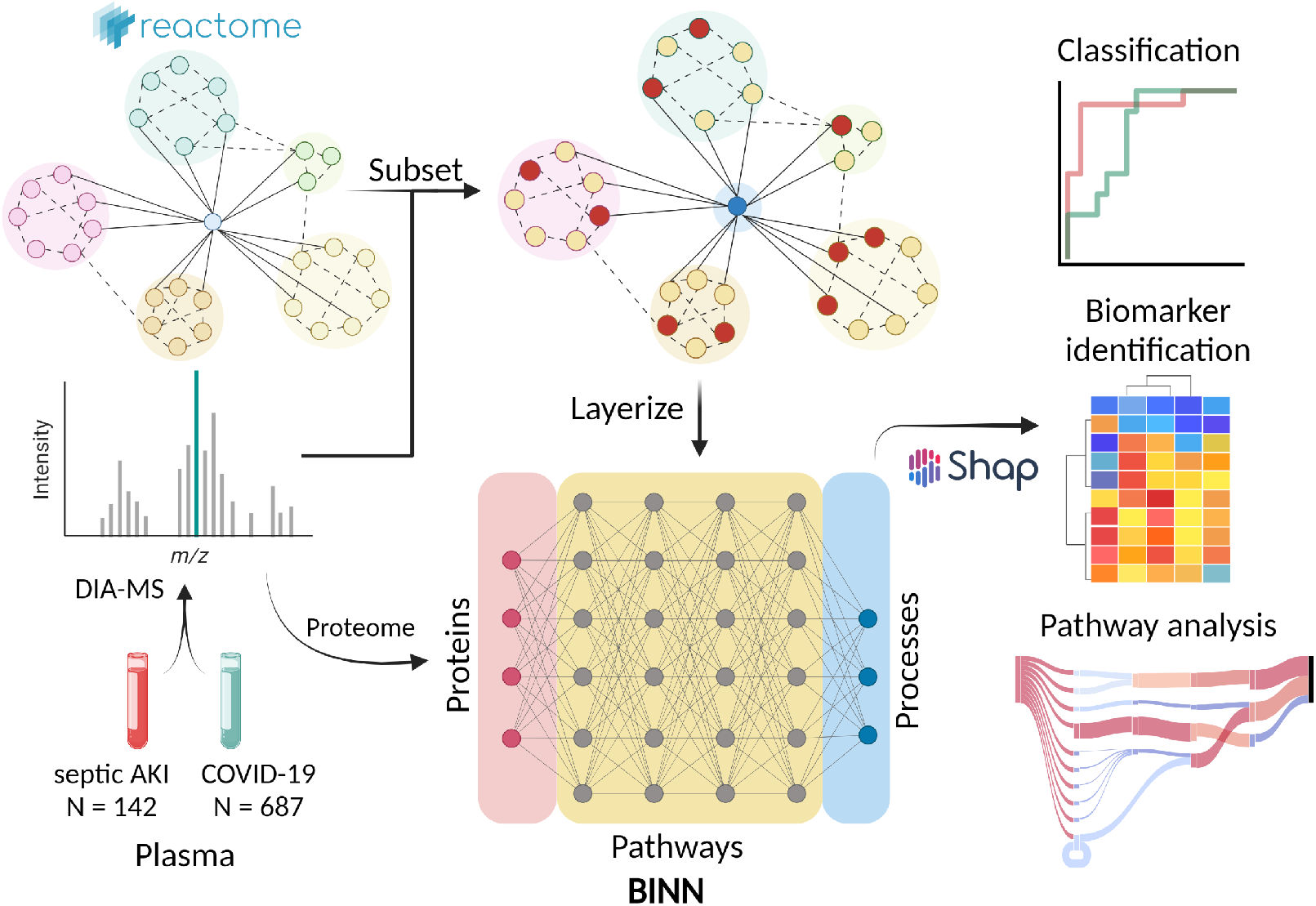
The complete workflow of analyzing proteomic data with biologically informed neural networks. The plasma proteome from patients suffering from septic AKI and COVID-19 were gathered and analyzed elsewhere [47, 29]. The data was downloaded and re-analyzed, resulting in datasets for the respective disease. The BINN is then generated by subsetting the Reactome pathway database [16] on the proteomic content of the respective dataset, and layerized to fit a sequential neural network-like structure. The protein quantities for each sample is used to train the BINNs to differentiate between two subphenotypes of COVID-19 and septic AKI respectively. Thereafter, the network is interpreted using SHAP and the resulting feature importance values allow for biomarker identification and pathway analysis. Created with BioRender.com.

### 2.1 Construction of biologically informed neural networks

As a starting point, proteomics plasma-data from patients suffering from septic AKI and COVID were analyzed to generate datasets for the respective disease. Septic AKI has previously been classified into two subphenotypes of varying severity by latent class analysis of various clinical markers [28]. 142 samples in the sepsis-dataset were stratified to one of the two subphenotypes, where 60 were classified as subphenotype 1 and 82 as subphenotype 2. Similarly, patients may suffer from varying degree of COVID-19, which has generated a scale defined by the World Health Organization to classify the severity of exhibited symptoms. According to this scale, patients requiring mechanical assistance for ventilation (WHO scale 6-7) are categorized as extremely severe, whereas patients able to breath by themselves as less severe, resulting in two subphenotypes of COVID-19. The COVID-dataset contained a total of 687 samples, where 406 were graded as very severe (WHO scale 6-7) and 281 as less severe (WHO scale <6). The proteomic content of the datasets differed, as 728 proteins were identified in the septic AKI dataset, as compared to the shallower proteome of the COVID-cohort containing 173 proteins.

The datasets were used in combination with the Reactome pathway database [16] to create and train BINNs. As mentioned, the Reactome database contains information about relationships of biological entities, such as molecules, pathways and high-level processes, and does not follow a sequential structure. The underlying graph is therefore subsetted and layerized to fit a sequential neural network-like structure, whereafter it is translated to a sparse neural network architecture, where nodes are annotated with a protein, biological pathway, or biological process - hence biologically informed neural networks. The proteomic content of a sample is passed to the input layer of the network, and the following layers map it to biological processes of increasing level of abstraction - finally ending up in high level processes such as *immune system, disease, metabolism etc*. The annotated and sparse nature of the network makes it well-suited for introspection and interpretation, as demonstrated by Elmarakeby et al. [25]. The algorithm which uses a graph and a subset of entities to create a sparse sequential neural network was generalized and implemented in the PyTorch-framework in Python, and is publicly available at GitHub: https://github.com/InfectionMedicineProteomics/BINN.

Networks for the respective disease were generated with four hidden layers each, and differed in architecture due to the discrepancy in the depth of the proteomes of the two datasets - the COVID-BINN being much smaller than the sepsis-BINN (supplementary 7). Due to their sparse nature, the resulting networks are small - containing trainable parameters in the thousands (sepsis-BINN: 6.7 k, COVID-BINN: 1.6 k trainable parameters between hidden layers), as compared to millions which is the case for most contemporary complex deep learning models. The BINNs were trained to identify the subphenotypes of septic AKI and COVID respectively, as outlined above.

### 2.2 Method comparison

To investigate whether machine learning methods were suitable for the stratification of septic AKI and COVID-19-subphenotypes, the BINNs were benchmarked against a support vector machine with radial-basis function kernel, k-nearest neighbour, a random forest, and two boosted trees (LightGBM and XGBoost). Evaluation was performed on the complete datasets using k-fold cross-validation. All machine learning methods achieved AUC-scores of >0.75, but the BINNs resulted in the best performances (ROC-AUC ^1^: 0.99±0.00 and 0.95±0.01, PR-AUC ^2^: 0.99±0.00 and 0.96±0.01) on the septic AKI and COVID-dataset respectively (figure 2A,B). Both BINNs achieved a high true positive and true negative rate (sepsis: 94 ± 2%, 100 ± 0%, COVID: 87 ± 2%, 92 ± 1%) (figure 2C). Additionally, both the sepsis and the COVID-BINN attained the highest precision and recall-rates out of all methods, achieving a precision of: 0.99±0.020, 0.87± 0.011, and recall of: 1.0 ± 0.0, 0.88 ± 0.022 respectively.

**Figure 2:**
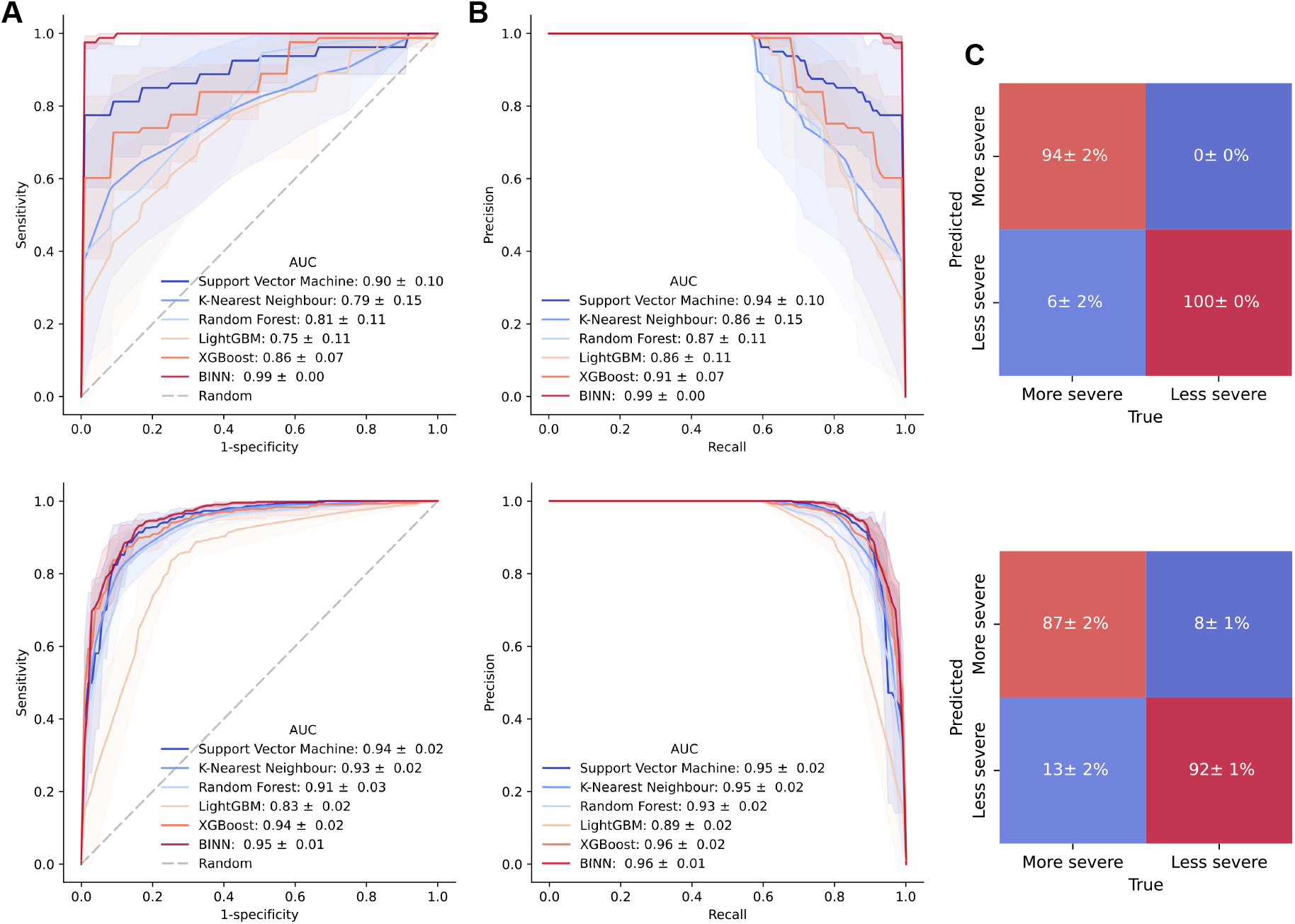
Performance of machine learning methods on the sepsis and COVID-datasets. The BINNs and five other machine learning models (support vector machine with radial-basis function kernel, k-nearest neighbour, random forest, LightGBM and XGBoost) were used to predict septic AKI and COVID-19 subphenotypes given the proteomic content of samples. The models were trained and evaluated using k-fold cross validation. **A)** The mean ROC-curve for the machine learning methods on the sepsis (upper) and COVID (lower). **B)** The mean PR-curve for the machine learning methods on the sepsis (upper) and COVID (lower). **C)** The confusion matrices for the respective BINNs (upper: sepsis, lower: COVID).

### 2.3 Interpretation

To identify which proteins, pathways and biological processes were important for the classifications, the trained BINNs were interpreted using SHAP [22]. SHAP is a feature attribution method which estimates the Shapley values (contribution) of each node in the network to the prediction. The node importance can be likened to how much worse predictions were to become after the removal of the said node. SHAP values were adjusted using the logarithm of the number of nodes in the reachable subgraph of a given node to account for the level of connectivity and to remove any biases associated with highly connected nodes (see supplementary methods and 10). The node importances of the complete networks were visualized in Sankey diagrams in figure 3. Nodes which were given a high SHAP value in the sepsis-BINN were largely related to metabolic processes, such as lipid metabolism and those related to PPAR-α [30], whereas the COVID-BINN places more importance on nodes related to the immune system and cell death. The emphasis on metabolic processes in the sepsis-BINN supports the view of sepsis as a condition with large systemic effects on metabolism, homeostasis and not solely the immune system [31, 32]. In the case for differentiating between COVID severities, processes relating to immune system (driven by innate immunity), metabolism of proteins, and programmed cell death, seemed to be the most important factors.

**Figure 3:**
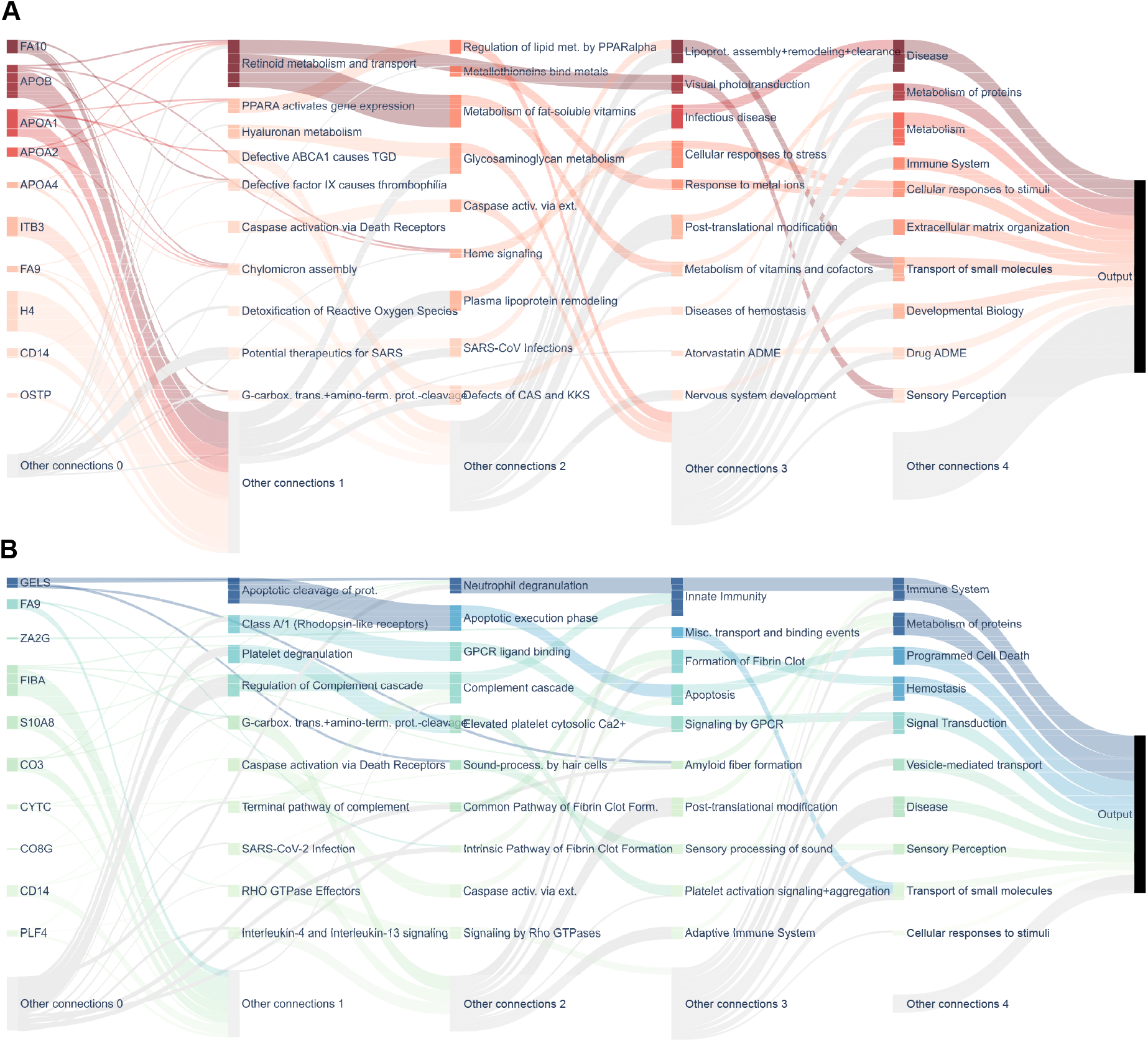
Sankey diagram visualization of node importance in the complete sepsis and COVID-BINNs. The importance for each node was calculated layer-wise using SHAP and reduced by the level of connectivity, and represented as the outgoing flow from the given node. Additionally, the nodes color reflects its relative importance, as darker nodes are more important in a given layer. The top 10 most important nodes in each layers are showcased and labeled, whereas the rest are gathered in the gray nodes at the bottom of the diagram (labeled ”Other connections”). Nodes that had no connection to the labeled nodes (i.e., both originated and targeted unlabeled nodes) were discarded for the sake of better visualization. **A) The sepsis-BINN.** Nodes related to metabolic processes, such as *lipoprotein assembly, remodeling and clearance* and *metabolism of vitamins and cofactors*, and disease, such as *infectious disease* are considered important in the sepsis-BINN. **B) The COVID-BINN.** In the COVID-BINN, processes related to *immunity, protein metabolism* and *programmed cell death* are dominating. This highlights a difference in the two networks, where subphenotypes of septic AKI are distinguishable by metabolic processes, whereas differentiating between COVID-19 subphenotypes relies more on pathways related to immunity.

#### 2.3.1 BINN-enhanced biomarker identification

The first layers of the BINNs contain the proteomic content, and to investigate whether proteins deemed important for the classification by the BINNs could be considered as potential biomarkers, the top-ranking proteins by SHAP value were subject to further investigation. For comparison, a measure of differential expression, the DE-score, was devised as a means of standardizing differential expression analysis. The DE-score is calculated by scaling the logarithmized fold change and *p-value* and computing their Pythagorean sum. Proteins which most significantly differ between two groups will therefore be given a high DE-score (equation 3, supplementary 8). Hierarchical clustering using Ward’s minimum variance method was performed on the protein quantities of the top 20 proteins identified by SHAP and by DE-score in both the sepsis and COVID-BINNs.

Several of the top-ranking proteins in the sepsis-BINN were known biomarkers for inflammation and have been documented to be altered during severe sepsis, such as CD14 [33, 34], FA10 [35], H4 [36], and OSTP [37], however, proteins related to metabolic processes, such as apolipoproteins (APOB, APOA1, APOA2 and APOA4) were also identified. Notably, these were not included in the top-ranking proteins by DE-score and wouldn’t be identified with classical differential expression analysis. Clustering on the proteins identified by SHAP resulted in a Rand-index of 0.765, outperforming the clustering on proteins ranked by DE-score which achieved a Rand-index of 0.716 (7.0% increase). Similarly, many of the most important proteins in the COVID-BINN have also been identified as biomarkers for the distinction between moderately and critically ill COVID-patients, such as GELS and ZA2G [38]. In the case for COVID, the differential expression analysis resulted similar proteins and results as the BINN, resulting in Rand-indexes of 0.645 and 0.663 respectively when performing hierarchical clustering (2.7% increase).

Markedly, the proteins with the highest SHAP value are not the most significantly differentially expressed or exhibit the highest fold change (see supplementary 9). This suggests that some proteins are considered important because of which pathways they are connected to, or due to their co-regulation with other proteins, and would likely have been discarded in typical analyses. Naturally, the proteins selected by DE-score differed in relative abundance, although the interpretable machine learning-centered method outperformed differential expression analysis in finding proteins which clustered to the subphenotypes. Clustermaps and plots showcasing the relative abundance of the identified proteins by SHAP can be seen in figure 4, and similar plots in the case of differential expression analysis can be seen in supplementary 6.

**Figure 4:**
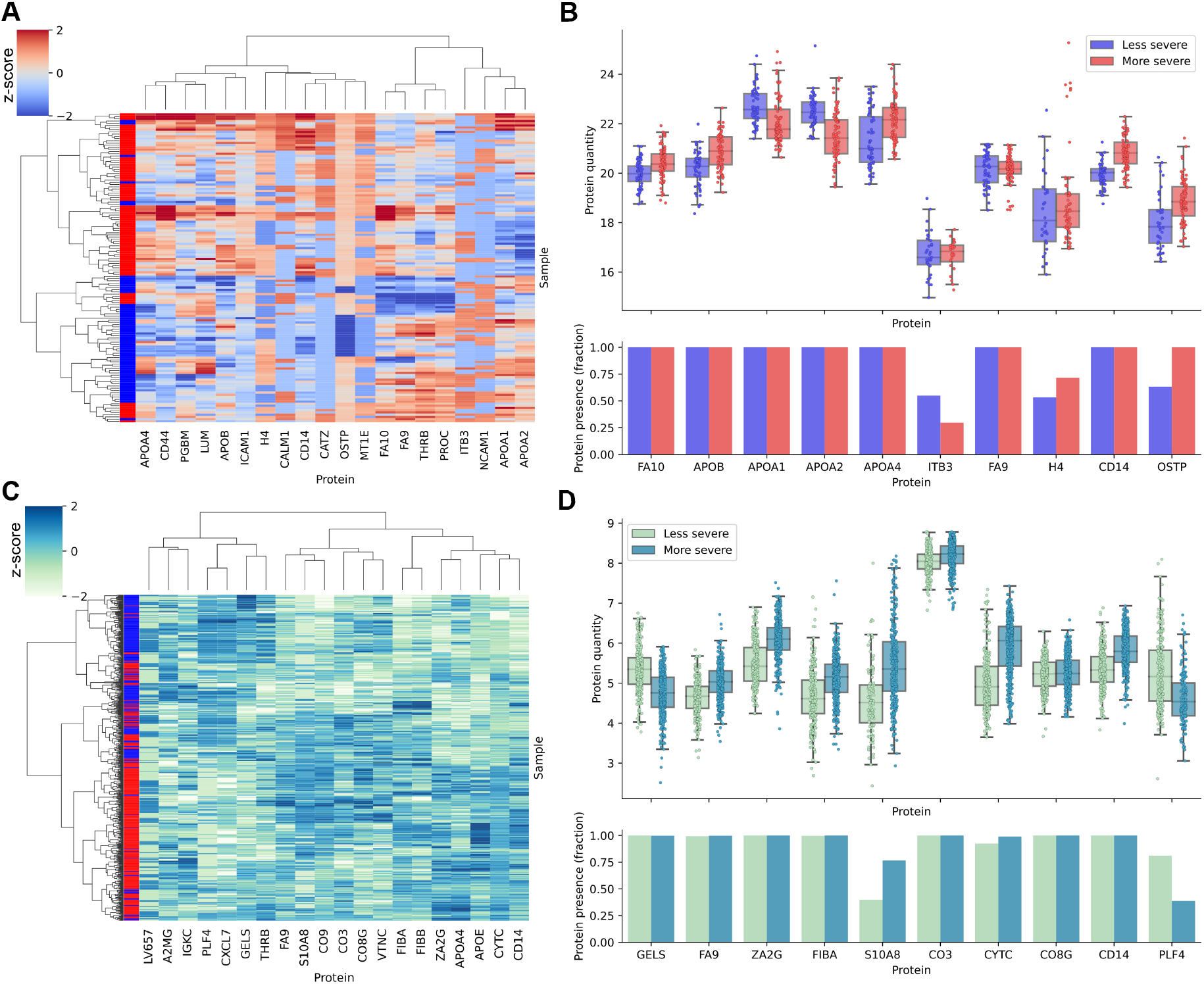
Clustering on the proteins with the highest SHAP values in the sepsis and COVID-datasets. The most important proteins as determined by the BINNs were selected and subject to hierarchical clustering. **A)** A clustermap showcasing the clustering based on the scaled protein abundances of the top 20 most important proteins in the sepsis-BINN. The left-most column shows the subphenotype classification (subphenotype 2: red, subphenotype 1: blue). Clustering was performed using Wards minimum variance method and Euclidean distances. The Rand-index for the clustering was 0.765. **B)** The upper panel shows the protein quantity for the 10 most important proteins. The lower panel shows which fraction of samples identified the given protein. **C)** Same as **A** but on the COVID-dataset. The Rand-index for the clustering was 0.663. **D)** same as **B** but for the COVID-dataset.

#### 2.3.2 BINN-enhanced pathway analysis

Since pathways and processes are integrated into the structure of the BINNs, a subset of pathways may be extracted from the graph underlying the BINN for pathway analysis. One may investigate pathways originating from a certain protein or pathway to see which pathways the node influences, and in turn, which it is influenced by. As mentioned, CD14 was identified as one of the most important proteins in the sepsis-BINN, and have many known implications in the immune response in general, as well as specifically in the course of sepsis [33, 34]. In figure 5A, CD14 has therefore been selected in the sepsis-BINN and the downstream pathways and processed visualized. In the network, CD14 funnels most of its importance through *caspase activation* and *TLR associated diseases*, and eventually to *disease, immune system* and *programmed cell death*.

**Figure 5:**
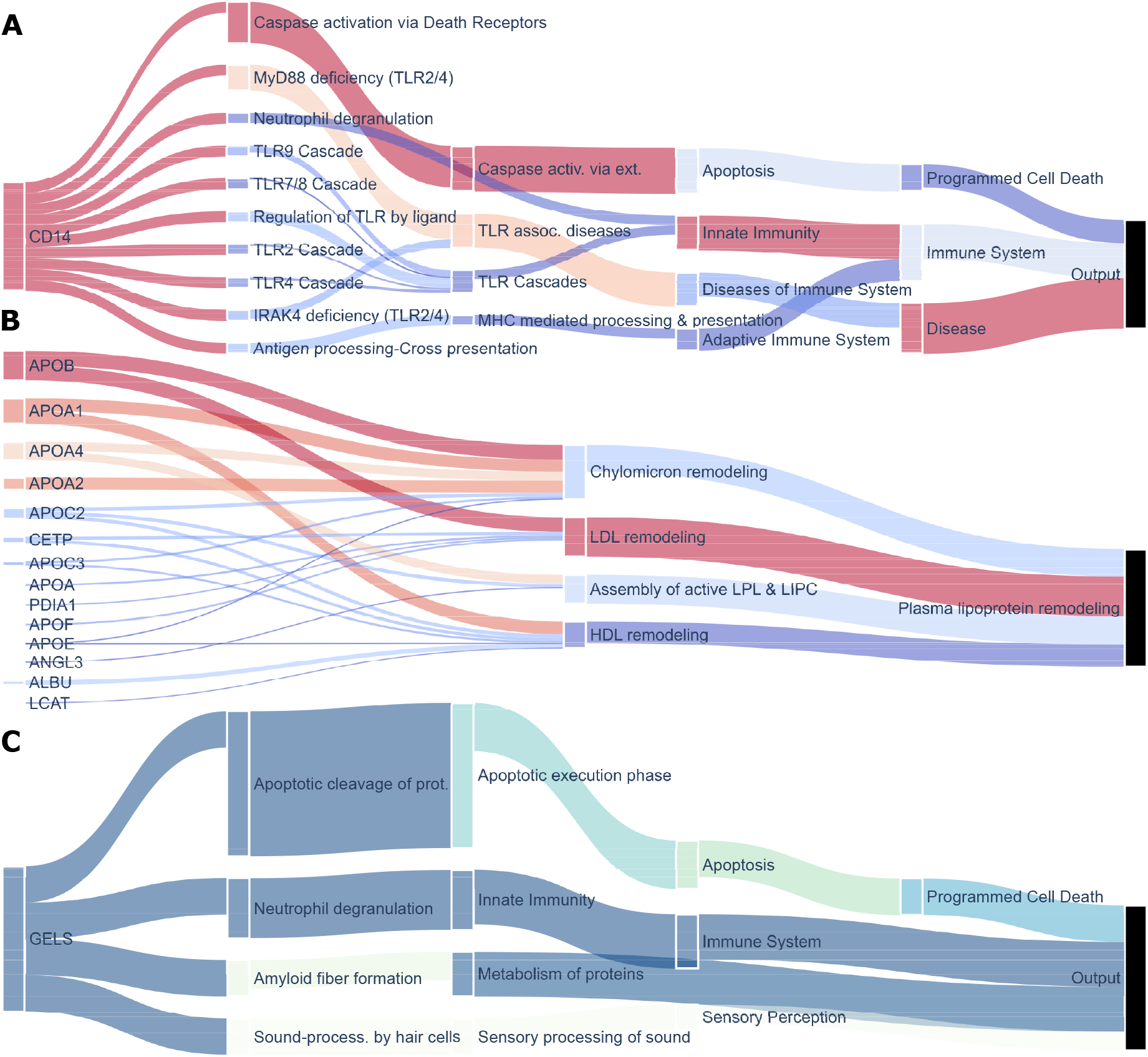
Custom pathway-analysis utilizing the interpreted BINNs. The graph underlying the interpreted BINNs can be extracted and subsetted for custom pathway analysis. One may look down-stream, i.e., recovering the subgraph originating from a specific protein. This showcases which pathways a protein will influence. One may also look up-stream, which instead showcases what proteins and pathways contribute to a specific process. **A)** The down-stream graph originating from CD14 in the sepsis-BINN. The most important contribution of CD14 is to *caspase activation via death receptors, MyD88 deficiency*, and subsequently, *disease* and *programmed cell death*. **B)** The up-stream graph originating from *plasma lipoprotein remodeling*. It’s most important contributor is *LDL remodeling, HDL remodeling* and four apolipoproteins: APOB, APOA1, APOA4 and APOA2. **C)** The down-stream graph originating from GELS in the COVID-BINN. GELS eventually connects to *programmed cell death, sensory perception, immune system, and metabolism of proteins* where *programmed cell death* and *immune system* are the most important high-level processes and *sensory perception* plays have little impact on the network.

Lipoproteins and lipoprotein metabolism are subject to major clinically relevant alterations during sepsis [32], and indeed many lipoproteins and related pathways and processes were identified in the sepsis-BINN, as described above. When inspecting the subgraph upstream from *plasma lipoprotein remodeling*, *LDL remodeling* and APOB, APOA1, APOA2 and APOA4 can be identified as the most important sub-process and proteins respectively (figure 5B).

GELS has been identified to play an important role in various physiological conditions, diseases and inflammatory processes [39], and was identified as one of the most important proteins in the COVID-BINN. After inspection of the subgraph originating from GELS, we identify that it contributes mostly to *apoptotic cleavage of proteins* and *neutrophil degranulation* - processes which eventually contribute to *programmed cell death* and the *immune system*. Both neutrophil degranulation [40] and programmed cell death [41] have been found to be pivotal in the course of severe COVID-19.

Pathway analysis plays a key role in understanding complex biological systems, and is naturally closely tied to proteomic content. To compare the integrated pathway analysis utilizing BINNs with common contemporary methods, pathway analysis with Metascape was performed [15]. This resulted in largely the same set of pathways ranking highly in both datasets, a majority of which are related to the inflammatory response (supplementary 11). Utilizing the interpretable nature of the BINNs and querying their underlying graphs allowed us to find important pathways and relationships which wouldn’t have been discovered using many contemporary methods, highlighting the advantages of the BINNs for custom pathway analysis.

## 3 Discussion

We present and apply a generalized workflow utilizing biologically informed neural networks (BINNs) and feature attribution methods for biomarker discovery and pathway analysis in a discovery proteomics setting. Although the BINNs are sparse and have few trainable parameters, they accurately predicted degrees of severity in both septic AKI and COVID-19 from the plasma proteome alone. The sparse and informed nature of the BINN incorporates biological pathways and processes into its architecture, tailoring it for introspection. Further, biological relationships which are typically overlooked in common methods are captured in the network, and therefore highly relevant information is incorporated into the analysis. Ultimately, this allows for a comprehensive analysis of proteomic data in a single unified method.

Interpreting the BINNs trained to predict different subphenotypes of septic AKI and COVID-19 identified several relevant biomarkers and pathways which wouldn’t have been identified using common methods of differential expression and pathway analysis. Furthermore, it highlighted key differences between the two diseases, as proteins and processes related to *metabolism* and *disease* were considered highly important in the sepsis-BINN, whereas the COVID-BINN favoured proteins and processes related to *immunity*.

Biomarker disovery in the context of BINNs is performed by calculating the feature importance of the initial layer of the network. Several of the most important proteins in the sepsis and COVID-BINNs were known biomarkers of the respective disease, however, they differed from the most differentially expressed proteins. Important proteins were not necessarily the most significantly differentially expressed (supplementary 9). The two methods may therefore be seen as complementary, and both may provide value to an analysis. Whereas differential expression analysis is guaranteed to provide proteins with a high fold change and low *p-value*, as this is the selection criteria, a BINN will provide the proteins which are important in a classification context when taking biological processes into account.

The major strength of BINNs lies in their embedding of pathway analysis into the architecture as the graph underlying the trained network can be extracted and subsetted to identify influential nodes in the subgraphs. This enables the investigation of downstream pathways from a given protein to understand the extent of its impact in the network. Similarly, the proteins and pathways upstream from a given node can be extracted to identify the extent of their influence. Comparatively, this provides a major improvement to how generic pathway analysis is commonly performed in proteomic research, where proteins associated with pathways are counted and the pathway with the most connections is considered the most relevant.

The performances of the BINNs relative to other machine learning methods differed between the datasets, as the performance of the COVID-BINN was comparable to other methods, while the sepsis-BINN outperformed other methods (figure 2). This is likely due to the combination of a higher dimensionality and smaller cohort-size of the sepsis dataset, suggesting that the BINNs are able to represent the feature space more accurately in complex datasets given fewer examples as compared to shallower learning-methods. Beyond performance, the varying proteome depths may also have implications on the conclusions drawn after interpreting the networks, as the underlying proteomes influences their architectures. Such effects should be kept in mind when comparing networks, as was done when identifying metabolic processes to be more important in the sepsis-BINN than the COVID-BINN.

It was found that hyperparameter-configuration had a significant influence on the distribution of importance in the network. Specifically, prolonged training durations resulted in a dependency on combinations of low abundance features such as antibodies, which although improved classification accuracy, are of less biological interest in this context. The BINN is highly dependent on the quality of the underlying graph, the dataset as well as the overlap with the dataset. Proteins which are not mapped to events in the Reactome pathway database are discarded in the analysis, and for small datasets the reduction in features may be detrimental. Unsupervised learning methods aimed at classifying nodes such as BIONIC [42] may be utilized to generate comprehensive networks encompassing a large majority of the proteome which could be used to generate BINNs. However, defining and annotating processes and pathways is still a manual and laborious task limiting the size of the BINNs. Our implementation is agnostic to the underlying graph and inputs used for the creation of the network, allowing for e.g., genomic or metabolomic data to be used in combination with different pathway repositories such as KEGG [43], GeneOntology [44, 45], or a custom curated set of pathways, to generate BINNs.

In summary, we demonstrate how BINNs can be trained, interpreted and visualized to provide a comprehensive analysis of proteomic datasets. The methodology behind the creation, analysis and visualization of interpreted BINNs has been generalized and is publicly available, opening up possibilities for further analyses and development in the realm of machine learning in discovery proteomics.

## 4 Method

### 4.1 Data

Blood-plasma from patients suffering from septic AKI and COVID-19 were gathered and analyzed elsewhere, whereafter the resulting proteomic datasets were uploaded to proteomeXchange [46] and made publicly available [47, 29]. The COVID-19 dataset consisting of the raw data matrix of quantified precursors and design matrix with patient annotations were downloaded from PRIDE (PXD025752)[48] and re-analyzed. The original study reports two cohorts from different hospitals whereof the samples gathered at Charité containing 687 samples was used here. The raw mass spectrometry files and spectral library for the septic AKI dataset were downloaded from PRIDE (PXD038394) and analyzed with an adapted version of the DIAnRT workflow[49] using GPS[47] for validation. Using OpenSwath (v. 2.6)[50], a first iteration of sub-optimal retention time alignment is performed followed by validation and refined retention time alignment using the highest scoring quantified precursors for each run. This process is repeated 3 times, with strict retention time alignment and mass correction on the final iteration followed by false discovery rate control at the global peptide and protein levels to generate a quantitative matrix.

### 4.2 Data processing

The septic AKI and COVID-19 datasets were processed in the same manner using the open source python package DPKS (https://github.com/InfectionMedicineProteomics/DPKS). The quantitative matrices were filtered to remove decoys and precursors that did not pass a 1% false discovery rate control at the global peptide and protein levels. Samples were then mean-normalized to remove any bias in the data and proteins were quantified using a python implementation of the relative quantification *iq*-algorithm [51]. Differential expression analysis was performed between each group of each dataset for proteins quantified in a minimum of 3 samples per group using linear models and multiple testing correction with DPKS. For input into the BINN, only proteins considered in the differential analyses were used as input, and missing values were imputed as 0.

### 4.3 BINN

The BINN was first introduced as P-NET by Elmarakeby et al. [25], and the architecture and methodology closely resemble the one they presented. Here, however, we introduce a generalized methodology as demonstrated in the context of proteomics analysis and present further applications of the informed network. The BINN is a sequential sparse feed-forward neural network which is generated using an underlying graph. The underlying graph used in this study is that of the Reactome pathway database [16] and contains information about relationships of biological entities, such as molecules, pathways and high-level processes. The graph is processed and layerized before it is translated into neural network in the PyTorch framework [52]. The generalized algorithm underlying the creation of a BINN from the Reactome pathway database was implemented as a Python-package:

1. Subset the Reactome pathways database (directed graph) using the union of proteins by adding the parental pathway, starting at the protein level, until the highest level of nodes is reached (nodes with out degree= 0).
2. Generate a network from the subsetted pathways and add an output node connected to the highest level of nodes. The number of output nodes correspond to the number of classes the network is set to predict.
3. Starting at the output node, traverse the network backwards for N layers If reaching a terminal node before N layers have been reached - add a copy of the previous node. This implies that the path depth ≤ N + 1.
4. Remove nodes which have not been traversed.
5. Finally, connect proteins to the final corresponding terminal nodes.

The constraints on the connectivity of the nodes renders the BINN tiny in comparison to most contemporary architectures. In this study, two networks were generated, originating from two different proteomics datasets: the first being analyzed blood-plasma from patients suffering from septic AKI, and the second from patients suffering from COVID-19. The sepsis and COVID-19 contained a total of 1203 and 174 proteins respectively. All proteins were not present in the minimum requirement of 3 samples per group or were not present in the Reactome database, reducing the final number of proteins to 728 (septic AKI) and 127 (COVID-19). The Reactome pathway database was downloaded 2022-07-14. When generating networks with 4 layers, this resulted in 6.7 thousand (septic AKI) and 1.6 thousand (COVID-19) trainable parameters between the hidden layers in total (supplementary 7).

The network is constructed so that the lowest level of entities exists in the inputlayer, and the level of abstraction increases as the network is traversed towards the output layer. The first layer (input layer) therefore contains the proteins, and are fed the scaled protein abundances. Thereafter follows the lower-level biological pathways from the Reactome database, such as *regulation of the complement cascade*. The final layer contains information about high level biological processes, such as *immune system, hemostasis, disease* and *metabolism*. The hidden linear layers are intercepted by *tanh*-activation layers, as well as dropout layers and batch normalization.

The BINN is interpreted using SHAP [22]. SHAP is a feature attribution method which computes the importance of a given feature to the outcome of the model. Similar to LIME [24], SHAP applies a linear relationship in its explanation model. Furthermore, the properties of the feature importance values are equivalent to the properties of the well-established Shapley values [53], which, in short, makes SHAP a feature attribution method which estimates Shapley values with a linear explanation model. SHAP provides a range of kernels which can be used for various models, one of which being the Deep SHAP-kernel, which similar to DeepLift [23] can be applied to deep learning models such as neural networks. In essence, Deep SHAP improves on the DeepLift algorithm, by approximating the conditional expectations using a set of *background samples*. Thereafter, the SHAP values can be approximated such that they sum to the difference of the expected model output (based on the set of background samples) and the current model output: *f*(*x*) – *E*(*f*(*x*)).

Problems arise if one wants the node importance to be meaningful for all layers in a sequential feed-forward neural network. This is because earlier nodes may completely rely on later nodes, and may not be important by themselves. For the node importance to reflect that which is both important in itself, and important in the context of the complete network, fully connected output layers are placed after each hidden layer, and the final prediction is computed as the average of all of the output layers. The output from each output layer is passed through a *σ*-activation function before being averaged.

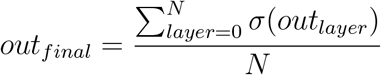

Nodes that are highly connected may be given an importance score which doesn’t reflect its biological importance, but is an artefact of the architecture. Elmarakeby et al. [25] used the graph informed function, *f*, to reduce bias that may be introduced by over-annotation of certain nodes:

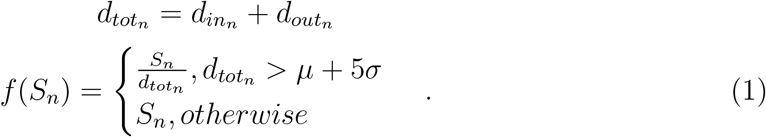

Where *d_in_n__* and *d_out_n__* are the in degree and out degree of a given node, n. To motivate the use of a bias reduction technique like this, we’d expect to see a correlation between the node degree and importance value. We suggest that a more general measure of node influence is the number of nodes in the complete subgraph defined by node n, *N_SG_n__*. The outgoing and incoming edges may be seen as a proxy for the size of *SG_n_*. The connections in a fully connected graph grows exponentially with the number of nodes, and *log*(*N_SG_n__*) may therefore be an appropriate reduction factor. Calculating the Pearson correlation coefficient for the mentioned graph informed measures and the SHAP value shows that the *N_SG_N__* and *log*(*N_SG_n__*) indeed are the graph informed functions that are most correlated with SHAP value, although this varies between layers (supplementary 10). The adjusted node importances may therefore be calculated by:

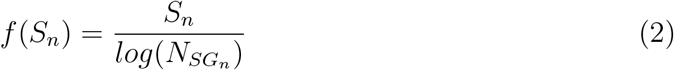

## 5 Training and evaluation

The generated datasets were scaled so that the distribution had a mean of 0 and variance of 1. The two generated BINNs were trained and evaluated on the respective dataset using k-fold cross-validation (*k* = 3) alongside five machine learning models (support vector machine with radial-basis function, K-nearest neighbor, random forest, LightGBM and XGBoost). Their performances were evaluated using the area under the receiver operator characteristic (ROC) curve and the area under the precision-recall (PR) curve. The area under the curves were averaged across cross-validation runs. The BINNs were trained until halted using early stopping and with similar hyperparemeter configurations. The learning rate was initiated at 0.001, and decreased adaptively if the validation loss plateaued. The networks seek to minimize the cross-validation error with an Adam-optimizer. A weight-decay (*L2*-penalty) of 0.001 is applied during training. Several measures were taken to mitigate the risk of over-fitting, such as the use of drop-out, as well as the adaptive learning rate, training times, and penalties mentioned above.

When generating models for interpretation, the BINNs were trained on the complete dataset, and never validated. In such cases, the adaptive learning rate is monitoring the training loss instead of the validation loss. Training was halted when training loss plateaued. The evaluation-time is dependent on the number of background samples used and it is often necessary to use a subset of the dataset as background, however, due to the relatively small number of samples in the dataset, the complete datasets were used to evaluate *E*(*f*(*x*)).

### 5.1 Biomarker evaluation and pathway analysis

Proteins, pathways and processes deemed important for the network during classification will be the ones that contribute greatly to correct predictions. The interpreted network can therefore be introspected and used for biomarker evaluation and pathway analysis. The proteins deemed important in the first layer can be extracted and compared to the ones that are differentially expressed. To get a quantitative measure of differential expression for a protein, *p*, the following expression was devised:

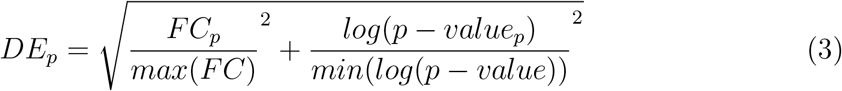

This normalizes the fold change (*FC*) and *log*(*p* – *value*) and calculates the Euclidean distance from *origo* (i.e., the Pythagorean sum). One can visualize this measure as the distance from *origo* in a volcano plot with a standardized scale on the x and y-axes (supplementary 8). The 20 proteins with the highest SHAP value and the highest DE-value were subject to hierarchical clustering using Ward’s minimum variance method.

One can subset the graph underlying the BINNs to extract subgraphs originating from, or incoming to, a certain node. The interpreted subgraph can be used for pathway analysis to gain further understanding of the dataset. We implemented three ways to subset the graph: downstream, upstream, and the combined downstream and upstream (complete subgraph). In a downstream subgraph, the pathways originating from a certain node is included, whereas in the case for an upstream subgraph, the nodes eventually reaching a certain node are included. A complete subgraph can be seen as the union of both the downstream and upstream subgraph.

**Figure 6:**
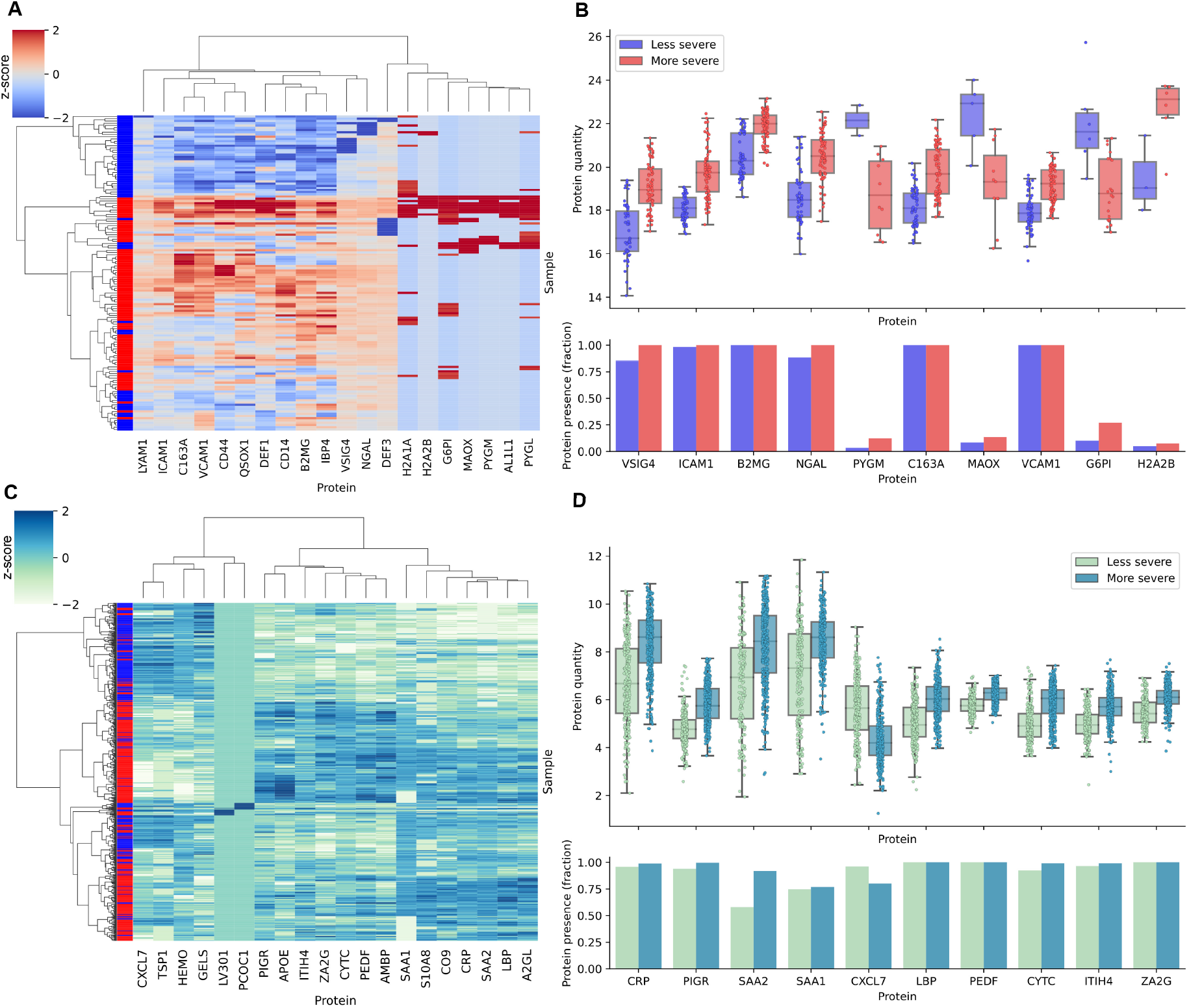
The most important proteins in the sepsis and COVID-BINNs by DE-score. The DE-score is defined in equation 3. The most important proteins as defined by DE-score were selected and subject to hierarchical clustering. **A)** A clustermap showcasing the clustering based on the scaled protein abundances of the top 20 proteins with the highest DE-score in the sepsis dataset. The left-most column shows the subphenotype classification (subphenotype 2: red, subphenotype 1: blue). Clustering was performed using Wards minimum variance method and Euclidean distances. The Rand-index for the clustering was 0.716. **B)** The upper panel shows the protein quantity for the 10 proteins with highest DE-score. The lower panel shows which fraction of samples identified the given protein. **C)** Same as **A** but on the COVID-dataset. The Rand-index for the clustering was 0.645. **D)** same as **B** but for the COVID-dataset.

## 6 Supplementary

**Figure 7:**
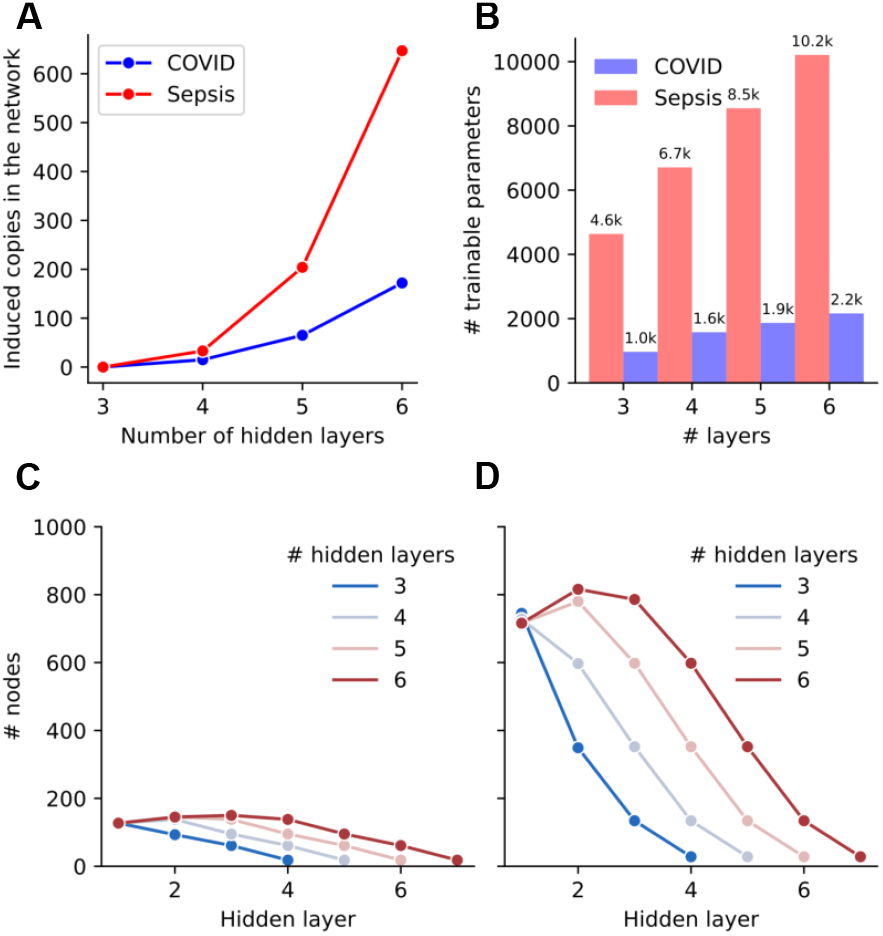
Features of the network architecture for the networks generated using the proteomic content of the sepsis and COVID-datasets. **A)** If the number of layers of the neural network surpasses the number of nodes in a given pathway, a copy is induced of the final node. Naturally, the number of copies increases as the number of hidden layers in the network increases. **B)** The number of trainable parameters in the network is in the thousands, and increases with the number of layers. The COVID-BINN has fewer trainable parameters than the sepsis-BINN, due to the lower number of proteins in the dataset. **C)** The number of nodes in the COVID-BINN when created with varying number of layers. **D)** The number of nodes in the sepsis-BINN when created with varying number of layers.

**Figure 8:**
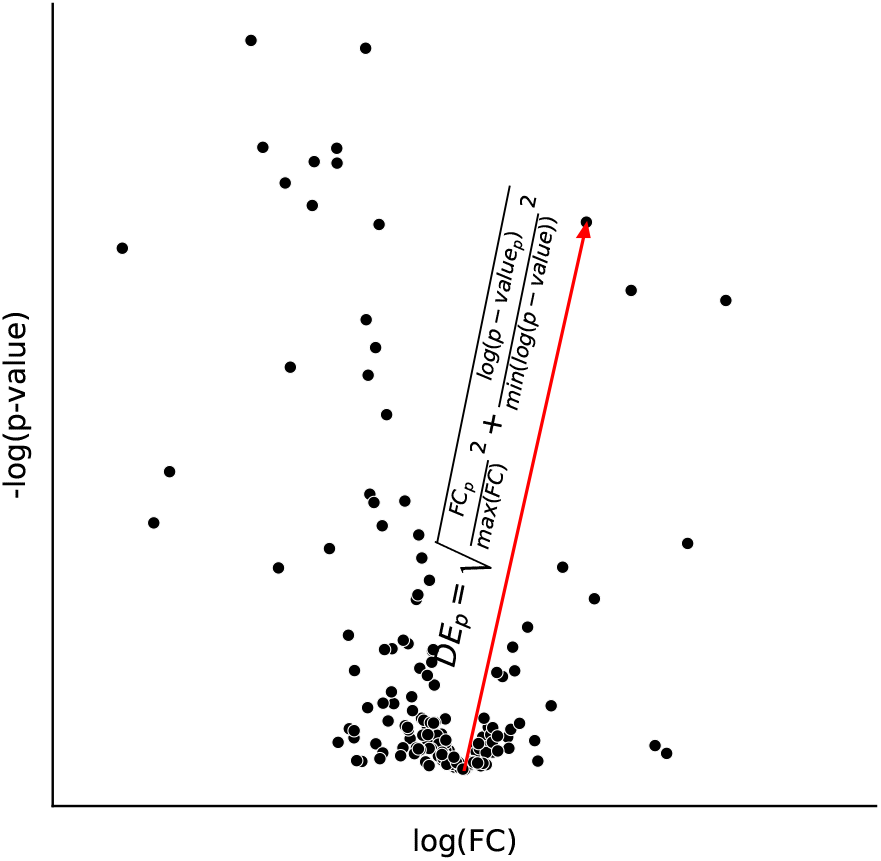
Visualization of the DE-score. To get a quantitative measure of level of differential expression, the DEscore was devised. The score can be seen as the Euclidean distance from *origo* to a protein in the volcano plot.

**Figure 9:**
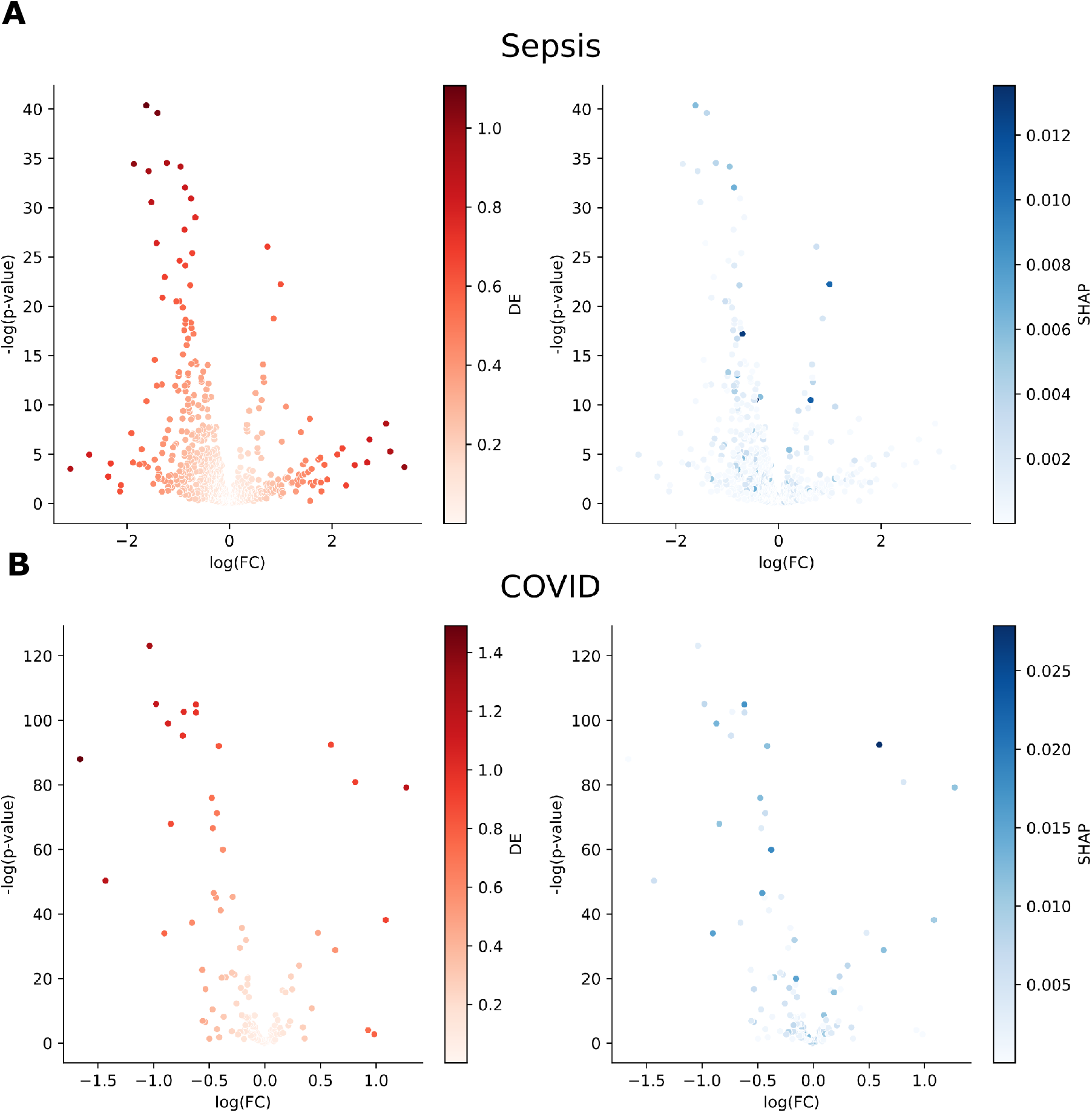
Volcano plots of the sepsis and COVID-dataset colored by DE and adjusted SHAP value. **A)** The volcano plots for the sepsis dataset. **B)** The volcano plots for the COVID-dataset. Coloring the volcano plots by DEscore (left) demonstrates how the most important proteins are selected by level of differential expression. When coloring by SHAP value it becomes apparent that SHAP value and level of differential expression are not highly correlated, as some proteins with low fold-change and high *p-value* are considered important for the BINNs.

**Figure 10:**
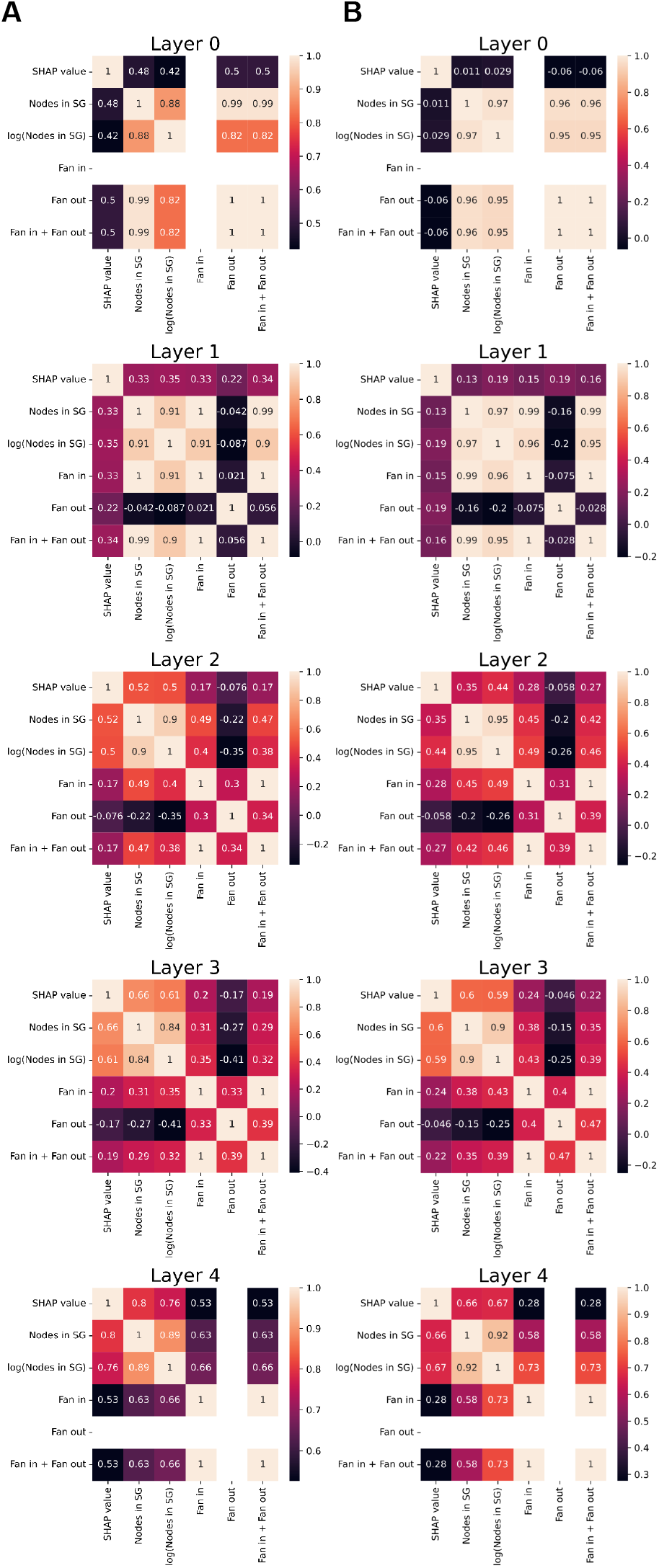
Correlation between graph features and SHAP value. Different graph attributes were correlated against each other and against the SHAP value. **A)** The correlation between graph features and SHAP value for the sepsis-BINN. **B)** The same but for the COVID-BINN. Nodes in a subgraph (*S_G_*, *log*(*S_G_*)) has the strongest correlation with SHAP value in most layers.

**Figure 11:**
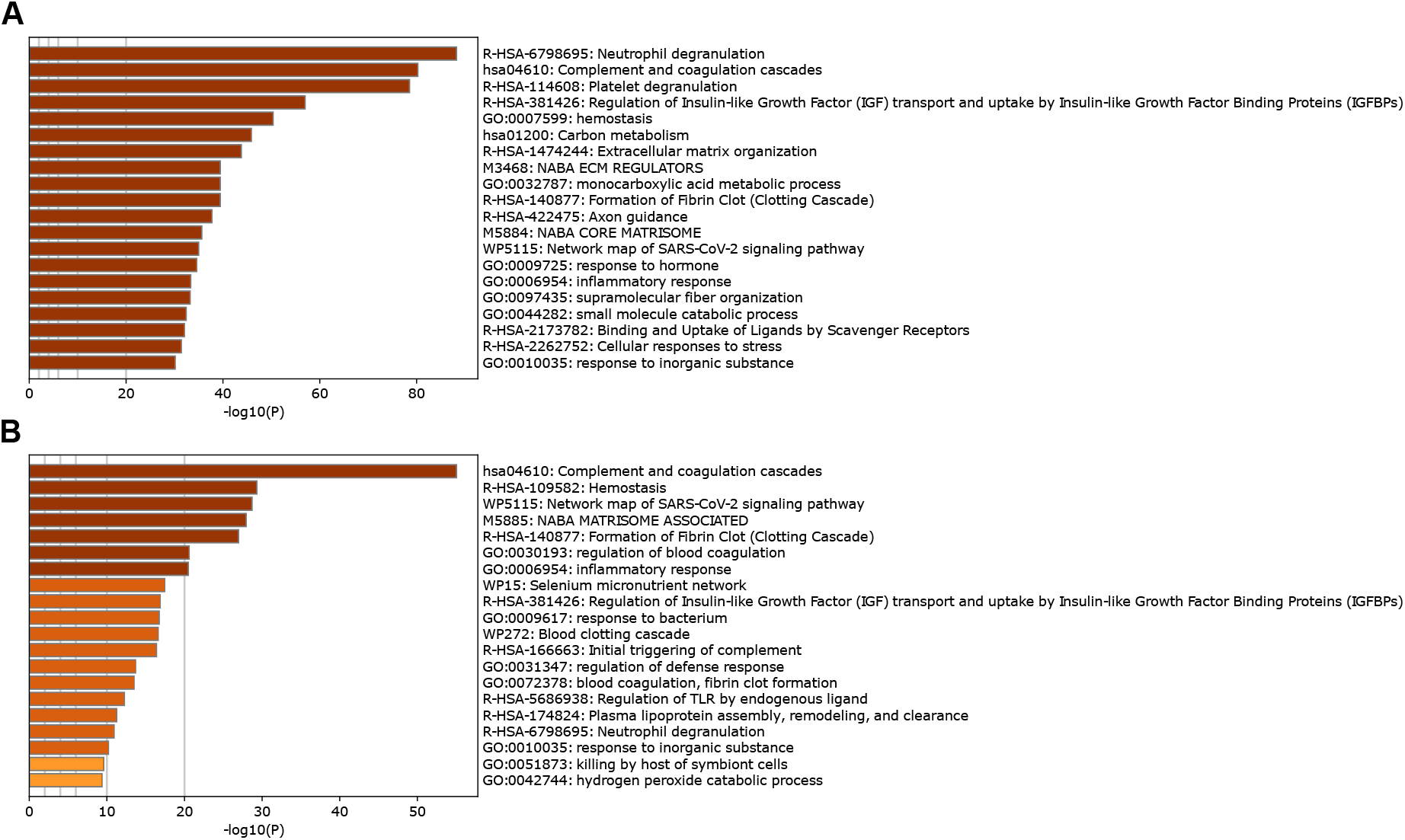
Pathway analysis of the sepsis and COVID-dataset using Metascape. The proteomic content was fed to Metascape [15] and the 20 pathways with the lowest *p-value* are shown. **A)** The top pathways identified in the sepsis-dataset. **B)** The top pathways identified in the COVID-dataset. There is a large overlap in the pathways and processes which are highlighted in the datasets, many of them relating to immunity and inflammation, such as the *complement cascade* and the *inflammatory response*.

## Footnotes

1 Area under the receiver operating characteristic curve

2 Area under the precision-recall curve

